# Neutralization of UK-variant VUI-202012/01 with COVAXIN vaccinated human serum

**DOI:** 10.1101/2021.01.26.426986

**Authors:** Gajanan N. Sapkal, Pragya D. Yadav, Raches Ella, Gururaj R. Deshpande, Rima R. Sahay, Nivedita Gupta, V Krishna Mohan, Priya Abraham, Samiran Panda, Balram Bhargava

**Author notes:** **Corresponding author**: Dr. Pragya D. Yadav, Scientist ‘E’ and Group Leader, Maximum Containment Facility, Indian Council of Medical Research-National Institute of Virology, Sus Road, Pashan, Pune, India Pin-411 021.

## Abstract

We performed the plaque reduction neutralization test (PRNT_50_) using sera collected from the recipients of BBV152/COVAXIN™ against hCoV-19/India/20203522 (UK-variant) and hCoV27 19/India/2020Q111 (heterologous strain). A comparable neutralization activity of the vaccinated individuals sera showed against UK-variant and the heterologous strain with similar efficiency, dispel the uncertainty of possible neutralization escape.

## Main text

The rapid surge in SARS-CoV-2 cases due to the Variant of Concern (VOC) 202012/01, also known as lineage B.1.1.7 or 20B/501Y.V1, in the United Kingdom (UK)^1^ raised concerns in several countries. Many of these countries had direct flights to and from UK and the variant was associated with high transmissibility. Identification of other variants from South Africa^2^ also initiated a global discussion on the challenges that these new variants could pose to the current vaccine candidates. The genome of the UK-variant has seventeen mutations, eight of which were found in spike receptor-binding domain (RBD) mediating the attachment of the virus to the angiotensin converting enzyme 2 (ACE2) receptor on the surface of human cells.^2^ Therefore, it appeared that the majority of the vaccine candidates, being either recombinant or specifically targeting the single epitope of original D614G ancestral spike sequence, might not be able to generate an efficient immune response against the new variants. Here, we successfully isolated and characterized the hCoV-19/India/20203522 SARS-CoV-2 (VOC) 202012/01 from UK-returnees in India with all signature mutations of the UK-variant.^3^ Earlier, we have reported development of an inactivated whole-virion SARS-CoV-2 vaccine BBV152 (COVAXIN™), which elicited remarkable neutralizing antibody response in phase I clinical trial against hCoV-19/India/2020770 (homologous), and two heterologous strains from the unclassified cluster namely hCoV-19/India/2020Q111 and hCoV-19/India/2020Q100.^4^ In phase II clinical trial, the vaccine candidate showed noteworthy results with plaque reduction neutralization test (PRNT_50_). The sero-conversion rate with neutralizing antibodies (NAb) following vaccination with BBV152 was 99.6%.^5^ Here, we present the NAb titers (PRNT_50_) of sera collected from vaccine recipients, who had received BBV152 vaccine-candidate in phase-II trial^4^ to underline the efficacy of BBV152 vaccine candidate against SARS-CoV-2 UK-variant (tested with sera of 38 human subjects) and one of the heterologous strains hCoV-19/India/2020Q111 (unclassified cluster) (tested with sera of 20 human subjects) using plaque reduction neutralization test as described earlier^6^. All the sera had equivalent NAb titers to hCoV-19/India/2020770 homologous strain and two heterologous strains with the characteristic N501Y substitution of the UK-variant; hCoV-19/India/20203522 (UK strain) as well as hCoV-19/India/2020Q111 (Figure 1 A and B). The median ratio of 50% neutralization of sera was found to be 0.8 when compared with hCoV-19/India/2020770 against mutant hCoV19/India/20203522 (UK strain), and 0.9 while compared with hCoV-19/India/2020Q111. Non parametric Kruskal-Wallis test for the comparison of the PRNT_50_ values from different groups revealed non-significant difference (p >0.05; Figure1). Andreano E *et al*. reported an escape of the UK-variant with E484K substitution, which was followed by an 11-amino-acid insertion in the NTD N5 loop (248aKTRNKSTSRRE248k) from high NAb in convalescent plasma, which was a serious concern pertaining to the potential efficacy of the vaccines.^7^ Our study evidently highlighted comparable neutralization activity of vaccinated individuals sera against variant as well as heterologous SARS-CoV-2 strains. Importantly, sera from the vaccine recipients could neutralise the UK-variant strains discounting the uncertainty around potential escape. It was reassuring from the PRNT_50_ data generated in our laboratory that the indigenous BBV152/ COVAXIN™, following its roll out in vaccination program, could be expected to work against the new UK-variant. It is unlikely that the mutation 501Y would be able to dampen the potential benefits of the vaccine in concern.

**Figure 1:**
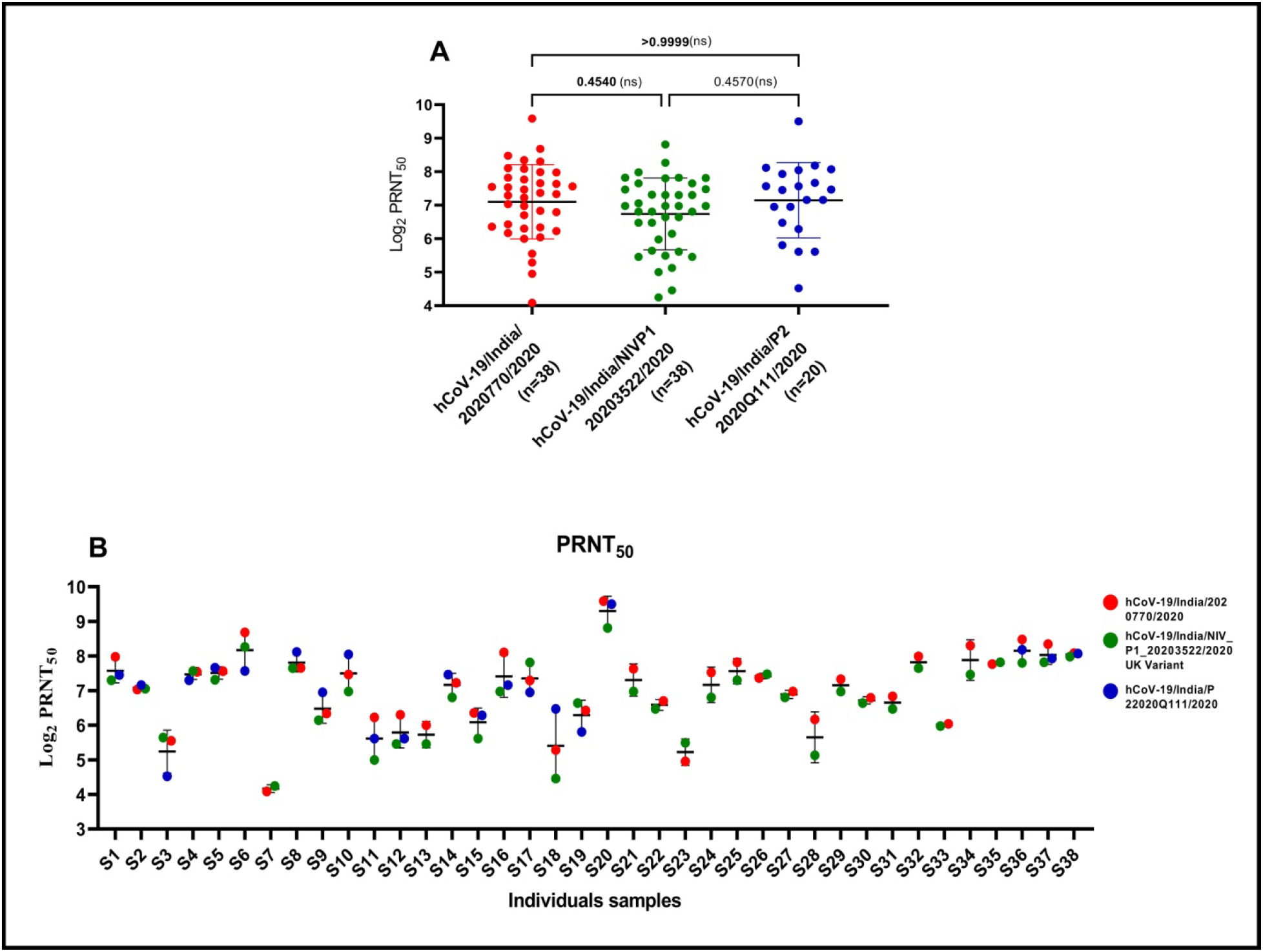
**(A) Neutralising antibody response of BBV152/COVAXIN™ vaccinated sera against SARS CoV-2 strains:** Neutralizing antibody titers (PRNT_50_) of vaccine’s sera against hCoV-19/India/2020770 (homologous), hCoV-19/India/20203522 (heterologous UK strain) and hCoV-19/India/2020Q111 (heterologous unclassified cluster). The bar represents the geometric mean and standard deviation of the respective group titers. Non-parametric Kruskal-Wallis test was used for the comparison of the PRNT_50_ values from different groups. The p-values above 0•05 were considered non-significant and are marked on the figure (ns-non-significant). **(B) Comparison of PRNT_50_ value of each vaccine recipient’s sera with three strains of SARS CoV-2:** Neutralizing antibody titers (PRNT_50_) of each vaccinees’ sera against hCoV-19/India/2020770 (homologous), hCoV-19/India/20203522 (heterologous UK strain) and hCoV-19/India/2020Q111 (heterologous unclassified cluster).The bar represents the mean and standard deviation of the respective sera.

## Acknowledgement

We thank the scientific staff of Maximum Containment Facility, ICMR-NIV, Pune Dr. Dimpal A. Nyayanit, Scientist C and Dr. Deepak Y. Patil, Scientist-B for providing excellent support. Also, authors are thankful to Darpan Phagiwala, Technician C, Prasad Gomade, Data Entry Operator, Diagnostic Virology Group, ICMR-NIV, Pune for excellent technical support.

## Data availability

The data that support the findings of this study are available from the corresponding authors upon reasonable request.

## Author Contributions

PDY and GNS contributed to study design, data collection, data analysis, interpretation, writing and critical review. GRD and RE contributed to data analysis and interpretation, writing and critical review. RRS, NG, VKM and SP contributed to data collection, writing and critical review. PA and BB contributed to writing and critical review of the manuscript.

## References

1. Kirby T. New variant of SARS-CoV-2 in UK causes surge of COVID-19. Lancet Respir Med 2021. doi: https://doi.org/10.1016/S2213-2600(21)00005-9

2. Starr TN, Greaney AJ, Hilton SK, et al. Deep mutational scanning of SARS-CoV-2 receptor binding domain reveals constraints on folding and ACE2 binding. Cell. 2020; 182 (5):1295–310.

3. Times of India. In a 1st, NIV-Pune isolates, cultures Covid's UK variant. https://timesofindia.indiatimes.com/india/in-a-1st-niv-pune-isolates-cultures-covids-uk-variant/articleshow/80078249.cms. (26 January 2021, date last accessed)

4. Ella R, Vadrevu KM, Jogdand H, et al. A Phase 1: Safety and Immunogenicity Trial of an Inactivated SARS-CoV-2 Vaccine-BBV152. medRxiv 2020. doi: https://doi.org/10.1101/2020.12.11.20210419

5. Ella R, Reddy S, Jogdand H, et al. Safety and immunogenicity clinical trial of an inactivated SARS-CoV-2 vaccine, BBV152 (a phase 2, double-blind, randomised controlled trial) and the persistence of immune responses from a phase 1 follow-up report. medRxiv 2020. doi: https://doi.org/10.1101/2020.12.21.20248643

6. Deshpande GR, Sapkal GN, Tilekar BN, et al. Neutralizing antibody responses to SARS-CoV-2 in COVID-19 patients. Indian J Med Res 2020; 152 (1): 82–87.

7. Andreano E, Piccini G, Licastro D, et al. SARS-CoV-2 escape in vitro from a highly neutralizing COVID-19 convalescent plasma. bioRxiv 2020. doi: 10.1101/2020.12.28.424451

